# Plant Prionome maps reveal specific roles of prion-like proteins in stress and memory

**DOI:** 10.1101/2020.09.25.311993

**Authors:** Sampurna Garai, Citu, Jyotsna Pandey, Sneh L. Singla-Pareek, Sudhir K. Sopory, Charanpreet Kaur, Gitanjali Yadav

**Affiliations:** Plant Stress Biology, International Centre for Genetic Engineering and Biotechnology (ICGEB), New Delhi, India 110067; Computational Biology Laboratory, National Institute of Plant Genome Research (NIPGR), New Delhi 110067; Stress Physiology and Molecular Biology Laboratory, School of Life Sciences, Jawaharlal Nehru University, New Delhi, 110067; Dept of Plant Sciences, University of Cambridge, U.K.

**Keywords:** Complex Network Analysis, *Oryza sativa*, Plant Amyloids, Prion-like Domains, Stress Biology, Stress Memory, Retrotransposons, Transcriptomics, Transposons

## Abstract

Prions can be considered as molecular memory devices, generating reproducible memory of a conformational change. Prion-like proteins (PrLPs) have been demonstrated to be present in plants, but their role in plant stress and memory remains largely unexplored. In this work, we report the widespread presence of PrLPs in plants through a comprehensive analysis of 39 genomes representing major taxonomic groups. We find diverse functional roles associated with plant ‘prionomes’. Investigation of the rice transcriptome further delineated the role of PrLPs in stress and developmental responses, leading us to explore whether and to what extent PrLPs may build stress memory. The rice prionome is significantly enriched for Transposons/Retrotransposons (Ts/RTRs), and we derived transcriptional regulatory inferences from diurnal gene expression revealing a complex regulatory network between PrLPs, transcription factors and genes known to be involved in stress priming, as well as transient and trans-generational plant memory. Overall, our data suggest that plant memory mechanisms may rely upon protein-based signals embedded in PrLPs, in addition to chromatin-based epigenetic signals and provides important insights into the anticipated role of prions in stress and memory.

## Introduction

Plant memory has emerged as one of the most fascinating fields of study in modern science, especially in view of the intricate mechanisms evolved by plants to survive under ever-changing unfavourable and adverse environmental conditions. One such phenomenon is related to the development of ‘stress memory’ in plants which can occur via ‘priming’, wherein a prior short exposure to stress, “primes” the plant for subsequent stress episodes by facilitating a faster and heightened response of resistance (Kinoshita and Seki, 2014; Crisp et al., 2016). Two major mechanisms have been reported to contribute towards ‘priming’ in plants; one being epigenetic in nature, and this is mediated by nucleosome remodelling via chromatin modification and changes in state of DNA methylation (Brzezinka et al., 2016). In contrast, the other mechanism is associated with heritable and self-perpetuating changes in the activity of proteins, mainly prions. Role of prions in plant memory is only beginning to emerge and these have been associated with diverse stress and memory processes in plants, including flowering time, as well as thermosensory responsiveness (Bailey et al., 2004; Shorter and Lindquist, 2005; Jung et al., 2020).

Prions are a special type of amyloid proteins, which can act as heritable elements in their aggregated state, constituting self-replicating entities with the ability to perpetuate and transmit over generations. They can switch from non-aggregated states to self-templating highly ordered aggregates and transmit this state to other homologous polypeptide sequences. This property allows them to confer stable changes in biological states that are of great interest in molecular and evolutionary biology (Newby and Lindquist, 2013). Prions have been extensively studied in several organisms (Alberti et al., 2009). In animals and fungi, these proteins are involved in roles ranging from devastating diseases in humans and mammals, to conferring beneficial heritable traits, such as cell survival under environmental stress conditions and memory (Prusiner, 1998; Pham et al., 2014; Sipe et al., 2016). In humans, diseases like Alzheimer’s, Parkinsons’s and Huntington’s syndrome have extensively been associated with amyloid aggregation with prion-like properties (Schwab et al., 2008). Prions are usually characterized by Prion-like domains (PrLDs), enriched in asparagine and glutamine (Q/N) residues (Dorsman et al., 2002; Fändrich and Dobson, 2002; Halfmann et al., 2011) along with glycine, serine and tyrosine (Kato et al., 2012). These PrLDs are not stringently infectious in nature as they are not transmitted between individuals and are believed to facilitate adaptation to the changing environmental conditions to which organisms are continuously exposed.

Plants are well known for their ability to sense cyclic changes occurring in their environment which may be compared to a memory process, wherein prion-like proteins (PrLPs) may provide a unique mode of biochemical memory through self-perpetuating changes in protein conformation and function. The screening of *A. thaliana* proteome has in fact, led to the identification of as many as 474 PrLPs (Chakrabortee et al., 2016), suggesting the need for a more comprehensive identification and general analysis of PrLPs in plants. Interestingly, plant flowering has been noted as a significant case for biological memory in Arabidopsis since its regulation involves memorizing and integrating previously encountered environmental conditions. Very recently, a prion-like domain (ELF3) in Arabidopsis, has been found to confer thermo-sensory responsiveness, a process that has previously been associated with epigenetic modulation, offering an opportunity for new studies that may bridge the two schools of thought in plant memory mechanisms (Jung et al., 2020).

In view of the above, we have addressed this knowledge gap through identification and dissection of the possible roles of PrLPs in the plant kingdom, which we collectively term as the ‘Prionome’. In all, we identified more than 4479 PrLPs in 39 plants, followed by an in-depth analysis of the density-enriched and well annotated rice prionome (201 PrLPs), in order to understand the possible correlation between physiological roles of PrLPs and stress priming. Interestingly, the rice prionome is significantly enriched in retro-transposon (RTR) like genes, predominantly localised in cytosol, mitochondria and the nucleus, showing significant expression under a variety of stress conditions. Further, we inferred transcriptional regulatory networks of PrLP genes and RTRs, using unweighted gene co-expression analysis. These networks revealed distinct clusters among rice PrLPs, which have evolved to form intricate linkages with hub nodes represented by transcription factors, which in turn, are either positively or negatively regulating genes reported to be involved in mediating transient and transgenerational memory in the plant kingdom. There is very little information on RTR genes in the rice transcriptome, and yet we find some RTRs to act as hubs in both positive and negative PrLP regulatory networks, involved in distinct sub-networks with genes involved in stress and memory. Taken together, our work is an attempt to unify the well-known epigenetic and lesser-known protein-based plant memory mechanisms and paves the way to further unravel cross-talk between factors contributing to “stress imprinting” in plants.

## Results

### Chlorophytes possess highest prion density in the functionally diverse plant prionome

Figure 1A depicts the density distribution of PrLPs across each of the 39 plant species used in the analysis, including four chlorophytes, two bryophytes (*Marchantia* and *Physcomitrella*), one gymnosperm (Pine), an ancient angiosperm (*Amborella*), six monocots (grass family members), and 25 dicots representing 13 taxonomic families. Algae possess highest prion densities, even with a threshold prion selection score of 25, with exception of *Micromonas pusilla CCMP* which was found to have the lowest (0.0076) densities across all phyla. Overall, lower plant taxa possess higher PrLP densities, with *Oryza sativa* and *Arabidopsis thaliana* depicting greatest prion density among all higher plants.

**Figure 1:**
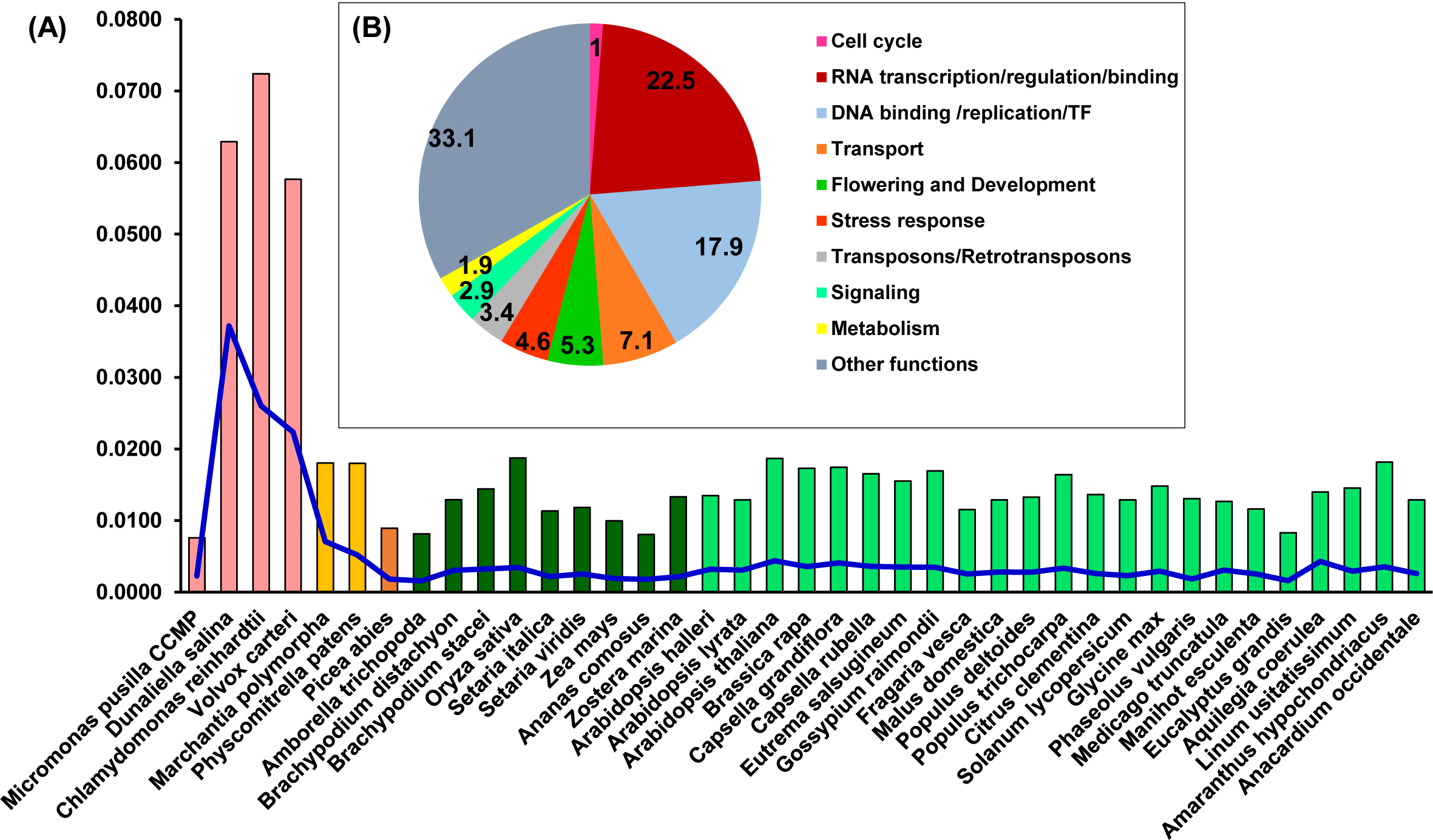
Distribution of PrLPs across the plant kingdom. (A) Bars show densities (PrLP count /Proteome size), blue lines show specific prion density at a threshold score of 25. (B) Ten Functional categories of PrLPs from 39 plants.

In all, we identified, 4479 PrLPs (above threshold core score value of 25) and these could be classified into ten functional categories depicted in Figure 1B. The representation was highest for the RNA binding/regulation/transcription (22.5%), category followed by DNA binding/replication/TF (17.9%) and transport (7.1%) (Supplemental Table 1). Notably, one of the top ten roles represented by PrLPs was flagged to be transposon (Ts/RTRs) related functions as can be seen in Figure 1B. These ten categories were further supported by gene ontology enrichment of plant PrLPs (Supplemental Table 2), with ‘nucleic acid binding’ function (GO:0003676) universally enriched across different phyla (Supplemental Figure 1A). Chlorophytes were specifically enriched in various functions related to DNA binding. Further, DNA/RNA-related functions like, transcription co-regulator activity (GO:0003712) and RNA binding (GO:0003723), were found to be co-enriched in *A. thaliana, P. abies* and *P. patens*. Flowering and development were overrepresented in various plant prionomes, along with proteins involved in the regulation of the reproductive processes (GO: 2000241). Plant prionomes were also enriched in other functions such as, aromatic/cyclic compound metabolic process (GO:1901362, GO:0046483, GO:0019438, GO:0006725 terms) in most species (Supplemental Figure 1A).

In context of biological processes, similar indications about overrepresentation of DNA-related functions could be seen, with DNA biosynthesis, metabolism and DNA regulation at transcriptional level, or gene expression, being the predominant biological PrLP functions across all phyla (Supplemental Figure 1B). Nitrogen metabolism related processes were found to be enriched in higher angiosperms (GO:0051171, GO:0034641 and GO:0044271).

Analysis of cellular location revealed enrichment of various nucleus-related roles of PrLPs (Supplemental Figure 1C). For example, PrLPs form part of the transcription factor TFIID complex (GO:0005669) in many prionomes, while RNA polymerase II holoenzyme (GO:0016591) and complex (GO:0030880) components could be associated with *P. deltoides, M. domestica, B. rapa and S. italica* prionomes.

### The Rice Prionome: Enrichment of Ts/RTRs

We identified 228 PrLPs (encoded by 201 genes) in the rice proteome (listed in Supplemental Table 3). The well annotated rice genome and availability of several gene expression datasets, offers a unique opportunity to delineate precise functional roles of PrLPs, and explore their involvement in stress, or in acclimation to stress memory.

Functional classification and GO enrichment of the rice prionome revealed numerous unique functional features (Figure 2 & Supplemental Figure 2). In terms of cellular distribution of rice PrLPs, we observed a preference for mitochondrial localisation followed closely by nuclear and cytosol-based localization, with instances of localization in more than one organelle (Figure 2C &D). Nuclear rice PrLPs were associated with DNA-related roles like transcription factors (TCP, auxin related), transcriptional activators (RSG and SWIRM domains), flowering regulation related (LEUNIG and FCA proteins), as well as the RNA binding FUS proteins and KH domain proteins (Figure 2E; Supplemental Figure 2). In addition, transport proteins such as ANTH/ENTH domains, as well as VHS/ GAT domain containing PrLPs were also predicted to be nuclear localised (Supplemental Table 3). In terms of biological processes, rice PrLPs were found to be enriched in regulation of flower development (GO:0009909), auxin activation pathways (GO:0009734) and G-protein coupled receptor signalling (GO:0007186) (Supplemental Figure 2). Molecular function enrichment highlighted ATP-dependant helicases (GO:0008026), lipid binding (GO:0008289) and involvement in nutrient reservoirs of cell (GO:0045735) (Supplemental Figure 2).

**Figure 2:**
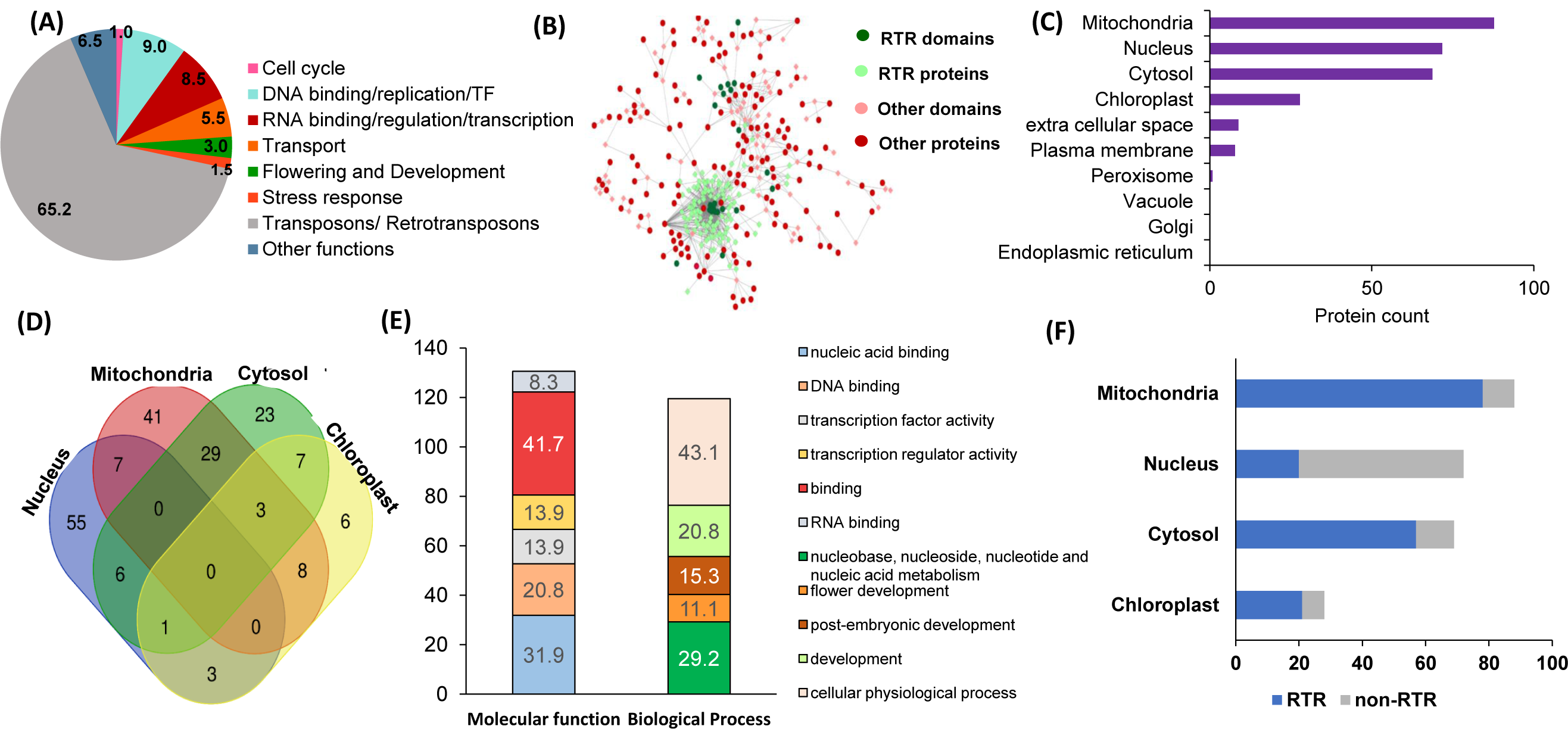
Rice prionome has preponderance of transposon/retrotransposon (Ts/RTR) type of PrLPs and participates in diverse functions of the cell with organellar enrichment of specific functions. (A) Functional classification (B) Domain categories including Transposons/Retrotransposons (Ts/RTRs) (Dark green) in Ts/RTRs candidates (light green), other domains (Red) in PrLPs (light pink diamonds). (C) Organelle-based distribution (D) Venn diagram depicting overlapping distribution of PrLPs in top four organelles. (C) Gene ontology-based classification of nucleus-localised rice PrLPs. (F) Distribution of Ts/RTRs in different organelles.

12 members in the rice prionome were identified as transcription factors, representing ARF, bZIP, C3H and NF-YB families, explored in more detail in later sections. Comparison of the rice prionome with the ten major functional categories described above, revealed the presence of an exceptionally large number of Ts/RTRs (Figure 2A & B, Supplemental Table 3). More than half (62.5%) of the identified PrLPs belong in this category (131 PrLPs). Interestingly, more than 80% of the cytosolic and mitochondrial rice prionome consists of Ts/RTRs (Figure 2F). RTRs were also detected in other plant prionomes, but nowhere as significantly as in case of rice, clearly indicating a unique case for *Oryza sativa*. Among other plants, chlorophyte *D. salina* (0.3%); monocots *B. distachyon* (1.2%), *S. italica* (1.3%) and *Z. mays* (3.9%); and dicots *A. thaliana* (3.8%), *G. raimondii* (1.3%) and *M. domestica* (3.9%) were found to harbour Ts/RTRs in their prionomes (Supplemental Table 1). We believe these numbers may not truly represent an absence of RTRs in other plants, but rather, a lack of sufficient or complete annotation about this important yet under-explored domain, and that future studies may reveal the existence of RTRs in other plant prionomes as well. For rice, the available annotation (RGAP version 7) has enabled a thorough investigation of the rice prionome, especially in terms of its unique enrichment for RTR/Ts domains.

### Rice prionome: Gene Expression Profiles

In order to gain deeper insights into potential prion-like roles of PrLPs, we analysed the development and tissue-specific expression profiles of 62 rice PrLPs, as described in methods (Figure 3). Expression levels of a stress-responsive N-rich protein, and a DAG protein-2 were found to be highest among all PrLPs across all stages of development (Supplemental Table 3). In contrast, RBD-FUS2, EXP3, EXP8 and EXP9 exhibited lowest expression among all rice PrLPs. However, few PrLPs showed significant variations during development such as, RNA-binding RBD-FUS1 protein, which was more abundant at the seedling and tillering stages and significantly downregulated during flowering stage. Further, transcript levels of ANTH/ENTH and RPA1C protein were increased at the flowering stage while those of BRO1, RSG activator and VHS and GAT1, were markedly downregulated during flowering. The TCP domain containing protein, SHR transcription factor, RRM1, ZFP1 and floral homeotic gene LEUNIG2 were upregulated in the stem elongation stage and downregulated at the heading stage. In coherence with the flowering and developmental functions, FCA has differential expression during vegetative and flowering stages (Figure 3A).

**Figure 3:**
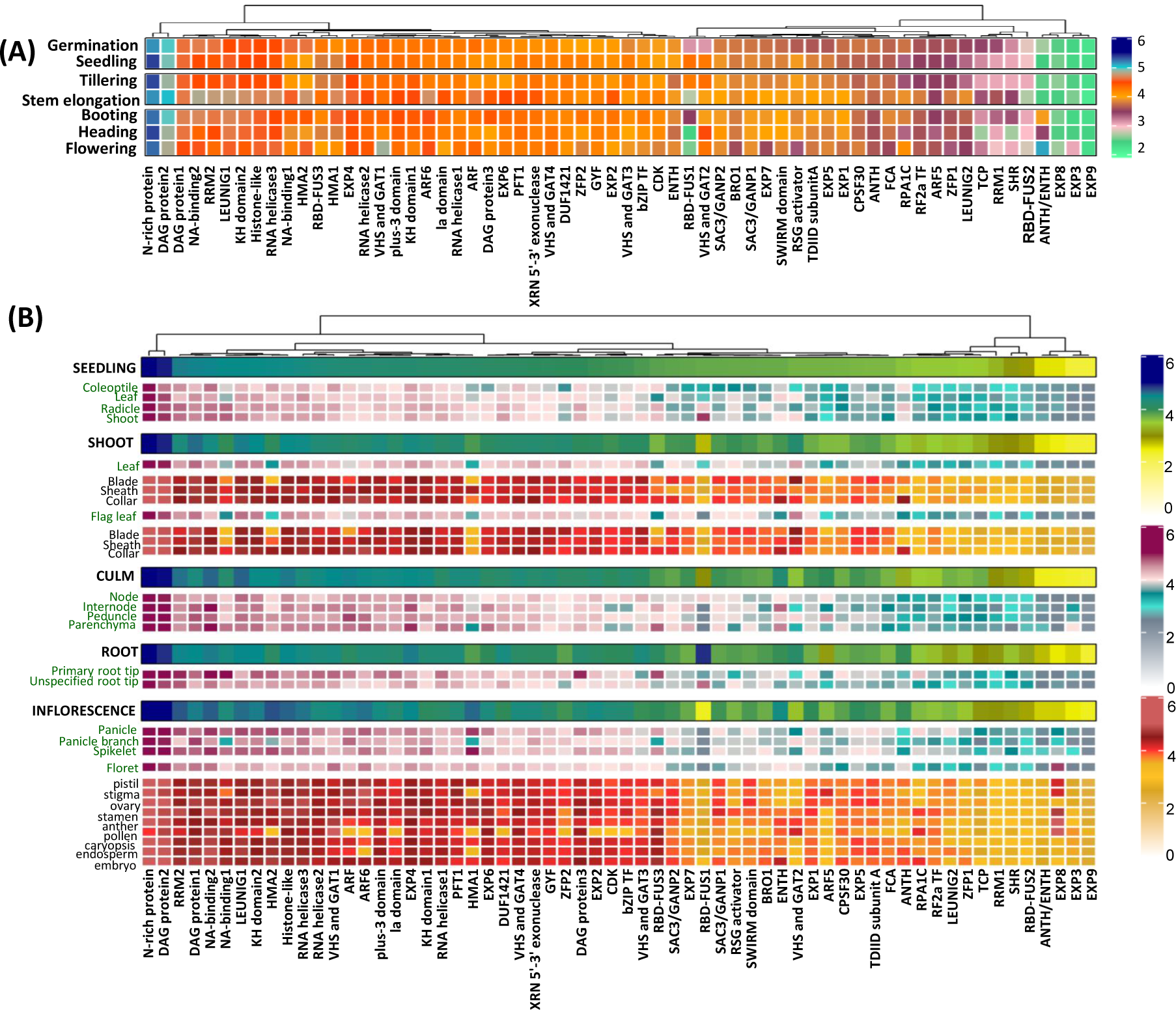
Differential Expression of rice prionome across **(A)** Developmental stages and **(B)** Data shown for 62 PrLPs obtained from Genevestigator. For full names and accession IDs refer to Supplemental Table 3.

Figure 3B depicts the rice prionome expression by tissue type. The most highly expressed group of PrLPs comprised of N-rich protein and DAG protein2 while SHR, RBD-FUS2, ANTH/ENTH and EXP3/8/9 constituted the least expressing groups across all tissues, as also observed across developmental stages. Differential expression of PrLPs was observed between male and female reproductive tissues, where ZFP2, ARF5, SWIRM domain, and SAC3/GANP1 showed much lesser expression levels in male tissues (stamen, anther and pollen) as compared to female tissues (pistil, stigma and ovary). Also, bZIP TF, CDK and ARF genes which were otherwise highly abundant in different organs showed lower expression in pollen whereas RPA1C showed opposite pattern of expression. Interestingly, stamen had upregulated expression for NA binding1 and RSG activator. The RBD-FUS1, however, showed greater expression in roots than shoots and inflorescence. In contrast, RRM2, NA-binding1, HMA1 and HMA2 are expressed more in the inflorescence. Further, VHS and GAT2 showed lowest expression in roots, particularly, in primary root tip and highest expression in the leaf blades. Importantly, root tips were particularly seen to express RBD-FUS3 and NA-binding proteins.

Overall, we noted that the plant prionome is extensively expressed in diverse tissues across developmental stages, and we then explored interaction between members of the prionome, as addressed in the next section.

### Rice Prionome Interaction network highlights essential cellular processes

Figure 4 depicts the protein-protein interaction network for the rice prionome, constructed as described in methods. Of the 201 PrLPs, 37 exhibit binary interactions with >1000 other members of rice proteome as evident from the core proteins (indicated in black circles) of the network. In order to functionally characterize the PrLP interactome, all 1263 interacting proteins were analyzed for pathway enrichment using KEGG database. Enrichment data was consistent with previous observations, showing DNA and RNA binding processes being the main biological roles of the rice PrLP interactome. In particular, functional clusters involved in ribosome and protein biogenesis, transcriptional machinery and its regulation, DNA replication/repair proteins and RNA surveillance proteins were predominant in the interactome (Figure 4). Since processing, transport and proteolysis of proteins involves a larger number of interactions, same could be seen from the interactome, with the largest cluster related to ribosome and protein biogenesis, represented by 886 proteins including 16 rice PrLPs. Similarly, flowering and development was also represented as a significant functional category in the interactome as observed in the rice prionome dataset alone. Mitochondrial biogenesis and autophagy, plant MAPK signaling pathway, and nucleotide and amino acid metabolism related functions were noted as other sub-networks.

**Figure 4:**
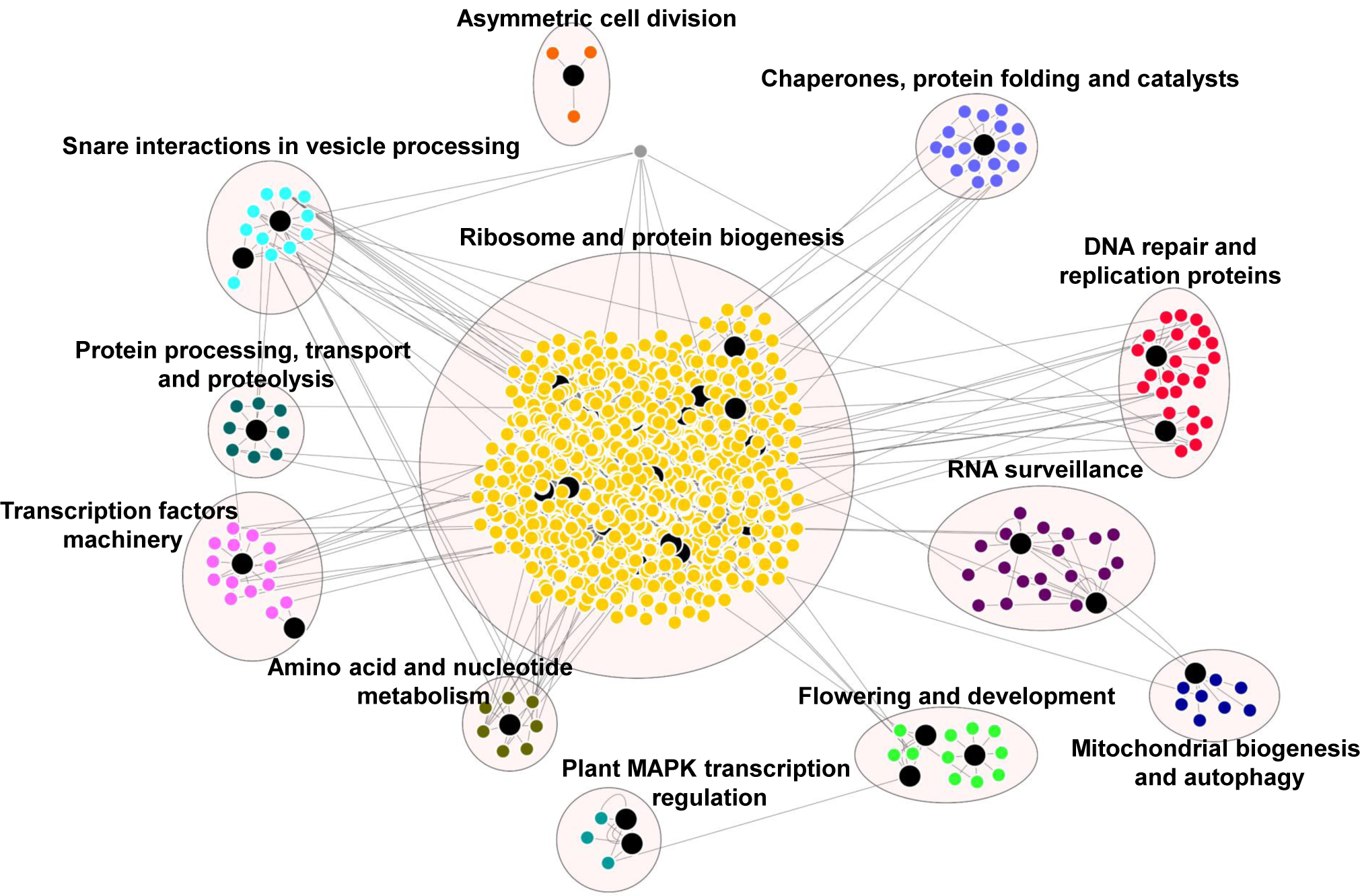
Rice prionome interaction network indicates role of PrLPs (large black nodes) in essential cellular processes. Different clusters depict proteins involved in the specific processes. Protein-protein interaction data was obtained from RiceNETDB and network was generated through MCA clustering in cytoscape.

### Role of PrLPs in Stress and Memory

We also investigated the response of the rice prionome to cold, heat, drought, salinity and biotic stresses, as shown in Figure 5. Assessment of the stress transcriptome map of the rice prionome identified specific PrLPs whose expression was significantly altered in response to stress conditions. For example, cold stress led to downregulation of RBD-FUS3, NA-binding1, DAG protein1, HMA1/2 and ZFP2 concomitant with upregulation of RBD-FUS1 and N-rich protein, which was found to be the most stress-responsive PrLP. Heat stress resulted in a nine-fold decrease in transcript levels of N-rich protein, while drought, salinity and *M. oryzae* infection also result in marked upregulation.

**Figure 5:**
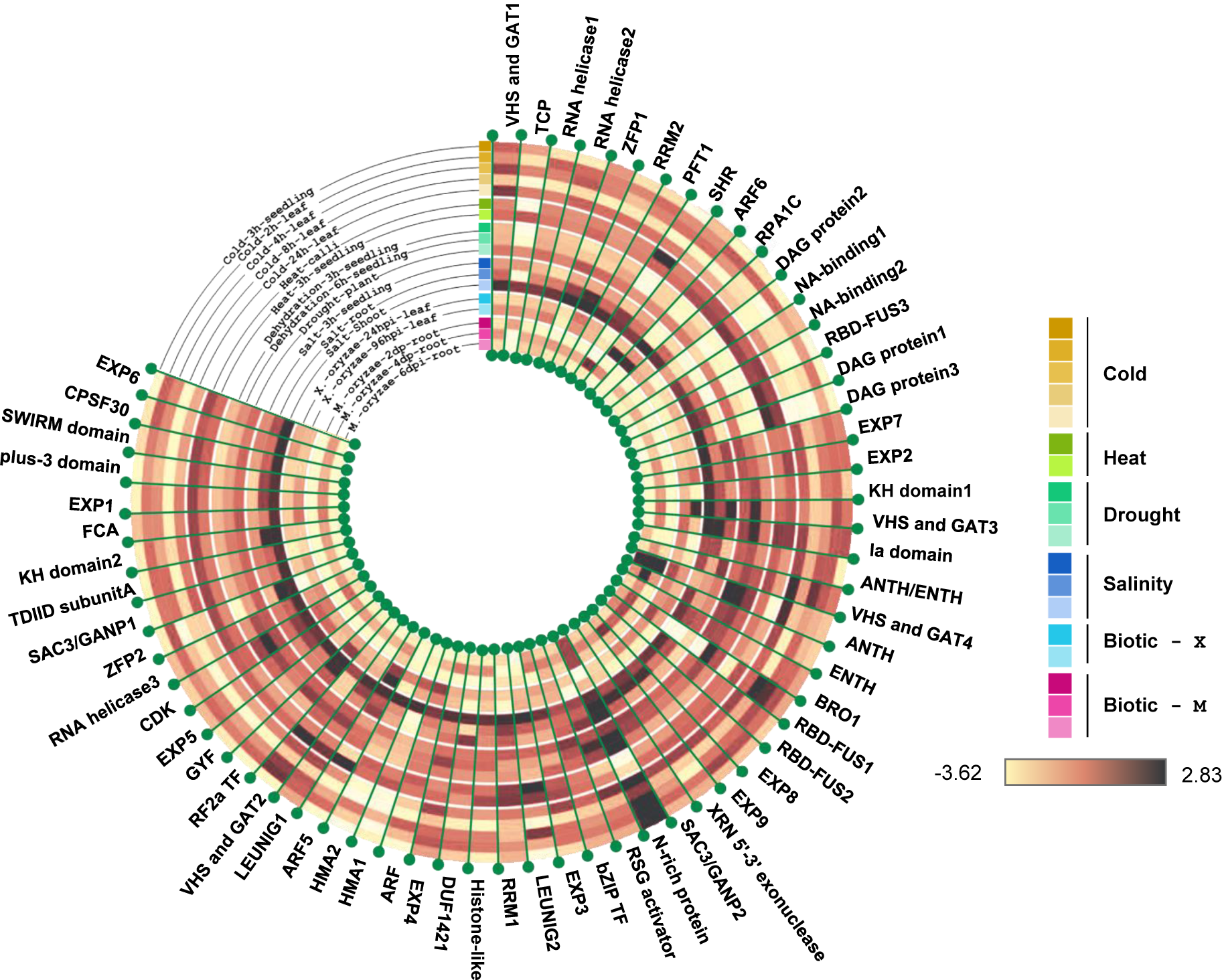
Rice prionome possesses members responsive to stress conditions (abiotic and biotic treatments). Tree constructed using R, shows log2 fold change in expression.

PrLPs encoding transcriptional corepressor LEUNIG1, auxin response factor ARF5, DAG protein1, VHS and GAT1 and SHR protein, are specifically increased in response to heat treatment, while the expression of TCP and ANTH genes is more than 4-fold downregulated under high temperature. Further, ANTH transcript levels are also drought-inducible. Notably, only two PrLPs namely, N-rich protein and VHS and GAT protein1, appear to be salt-inducible. Biotic stress responsiveness of the rice prionome was only observed for N-rich protein and ANTH. Overall, stress profiling of the rice prionome suggested differential regulation, catering to diverse processes from development to stress.

Interestingly, we found PrLPs to be involved in abiotic stress memory and stress recovery in several plants species, as depicted in Supplemental Figure 3. These range from cold stress in Arabidopsis (Zuther et al., 2019), hormonal stress priming in *M. domestica* (Supplemental Figure 3B) and so on. PrLP expression profiles in memory responses pertaining to recovery phase have been observed in *Populus spp*., in response to periodic and successively increasing drought, or chronic phase of combined drought-heat stress followed by one week of recovery phase (Supplemental Figure 3C). Likewise, heat stress showed a memory response among PrLPs in *Chlamydomonas* and *Arabidopsis* as well (Supplemental Figure 3 C&D). Interestingly, we could detect homologs for ten of these genes within the rice prionome, supporting the role of rice PrLPs in memory signals. These observations when combined with the large number of rice PrLPs found to be impacted by heat stress as mentioned in earlier sections, suggest an important role of the prionome in heat stress and memory. Notably, the very recent reports of heat shock proteins being important epigenetic mediators of transient as well as trans-generational memory, led us to explore cross-talk between reported epigenetic signals of memory and signals mediated by PrLPs. A full list of genes reported to be involved in plant stress or memory is provided in Supplemental Table 4.

### Transcriptional regulatory inferences from gene co-expression data

Gene co-expression data analysis was performed on rice PrLPs using the publicly available diurnal gene expression dataset, as described in Methods. Interestingly, the 66 PrLPs for which we found diurnal expression profiles (listed in Supplemental Table 5) included 11 of the 12 transcription factors in the rice prionome, as well as five Ts/RTSs, enabling a thorough investigation of the regulatory role of both TFs and RTRs in the prionome. The genes found to be significantly correlated with PrLPs, were used to (a) identify correlated clusters of genes, if any, among PrLPs, (b) capture the intersection between PrLPs, specially the Ts/RTRs, and genes known to be involved in stress or memory, and (c) identify transcription factors defining or influencing the rice prionome for insights into the PrLP master regulatory network.

Figure 6 depicts the correlation plot of the rice prionome, highlighting the significantly correlated PrLPs (at correlation coefficient cutoff of +-0.8 and P < 0.01). As can be seen in this Figure, two clusters of PrLPs are discernible, with about 15 genes in each cluster, and all 91 interactions are listed in Supplemental Table 6. Both clusters are significantly negatively correlated with each other, suggesting an antagonism in their roles/involvement, with one cluster including four TFs and two Ts/RTRs, while the other cluster having one TF and one Ts/RTR. Interestingly, the first cluster has several RNA helicases and exonucleases, as well as some of the PrLPs noted in earlier sections to be among the most highly expressed (DAG proteins) and those upregulated in floral tissue, both male (RSG) and female (ARF5 and ZFP2). In contrast, the second cluster, showing negative correlation with the first one, contains the BRO1 and RSG activators, whose expressions were earlier observed to be downregulated during flowering stages. Notably, this cluster also has the two LEUNIG proteins, well known to repress several floral homeotic genes in the floral meristems, required for proper differentiation of stamen and carpel structures in the flower (Franks et al., 2002; Sridhar et al., 2004). These patterns suggest a role for PrLP-mediated regulation among flowering genes. As can be seen in the inset, the first cluster also has several genes that were observed to be down-regulated in cold stress (RBD-FUS3, DAG, ZFP2) and upregulated during heat stress (ARF5, DAG1).

**Figure 6:**
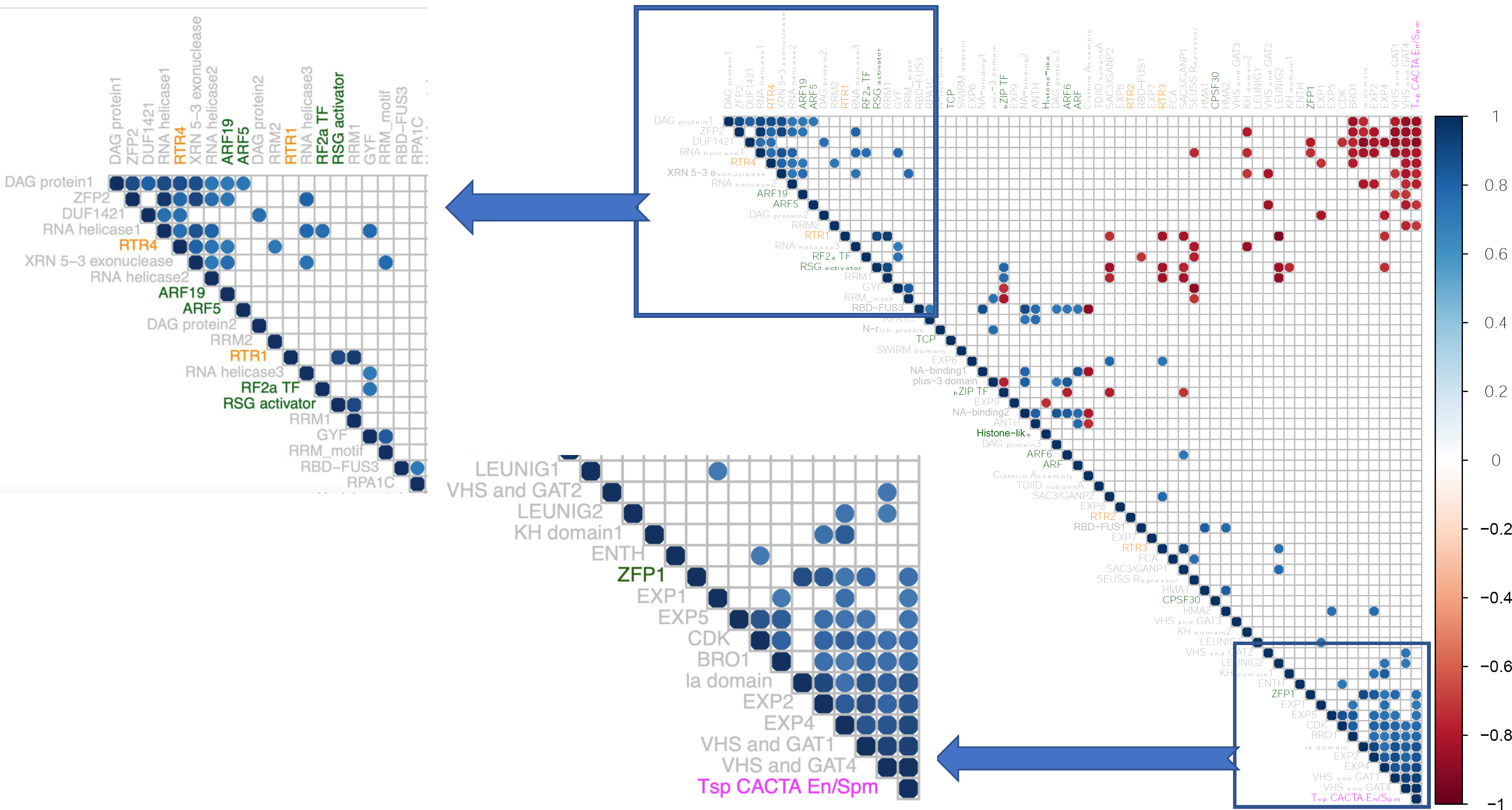
Correlation plot of 66 rice PrLPs (listed in Supplemental Table 5). Only significant correlations are shown (cut off +- 0.8 at P < 0.01). Rows/Columns ordered by First Principal component. Color of PrLP names depict TFs (Green), RTRs (orange) and Tsp (Magenta). Insets show two clusters, mutually antagonistic.

The composition of the above two PrLP clusters in diurnal co-expression data and the pattern of distribution of their respective transcription factors, corroborated by observations from condition-specific, tissue-based and developmental gene expression profiles, motivated us to derive regulatory inferences from co-expression data, for all TFs in the rice prionome.

As stated earlier, eleven of the 12 TFs in the rice prionome were found to have diurnal expression profiles and these 11 genes showed a significant positive correlation with 100 other TFs in rice over the entire day night cycle, as well as a significant inverse correlation (cor value < −0.8 at P < 0.01) with another 101 TFs, suggesting a master regulatory role for these 11 members of the rice prionome. To further ascertain a master regulatory role, we checked the upstream regions of all the positively and negatively correlated TFs for presence of cis-binding elements for the eleven PrLPs, resulting in the identification of 40 high fidelity rice TFs that had a strong positive or negative correlation with the rice prionome, in addition to containing the respective TF binding sites on their promoter sequences, and these have been added to Supplemental Table 6. These high fidelity TFs belong to the WRKY, Dof, C2H2, Myb-related, LBD, TCP, GATA, G2-like, TriHelix, SBP, RAV and NAC families, while eight of the 40 are PrLPs themselves, further supporting internal cross-talk and diverse regulatory roles of the rice prionome.

### Master Regulatory network reveals PrLP clusters in Memory acclimation

In order to visualise the patterns of interaction and regulation within and between the rice prionome and sets of other rice TFs or stress or memory genes (for which we had diurnal expression profiles), we constructed a transcriptional regulatory network using gene-co-expression data, and this was generated in a stepwise manner, as described in methods. This network enabled us to explore crosstalk between TFs and RTRs in the prionome, and the extent to which they may regulate or be controlled by other activators or repressors, especially in stress or memory acclimation.

The five Ts/RTRs in the rice prionome were found to be positively co-expressed with 91 TFs and negatively co-expressed with 77 distinct TFs. Of these pairs of co-expressing TF partners, we performed the same filtering as was done for 11 TFs above, to identify/retain only high fidelity TFs that have a known binding site on the respective Ts/RTR promoters (Supplemental Table 6). This was achieved by scanning the upstream regions of all co-expressing partners for the presence of cis-elements, leading to the retention of 22 true positive TFs, which were added to the rice prionome co-expression data, to generate a regulatory network. In the next step, the network was expanded by adding the 32 additional TFs that were identified earlier as high fidelity co-expressing partners of eleven TFs in the prionome. This network was then superimposed with available information on genes involved in stress or memory acclimation, by adding co-expressing partners of Ts/RTRs and TFs in the rice prionome that were (a) present in the rice stress interactome (Wierbowski et al., 2020) or (b) implicated in memory acclimation (Supplemental Table 4). The resulting master regulatory network is depicted in Figure 7, and the corresponding annotated edge list is provided as Supplemental Table 7.

**Figure 7:**
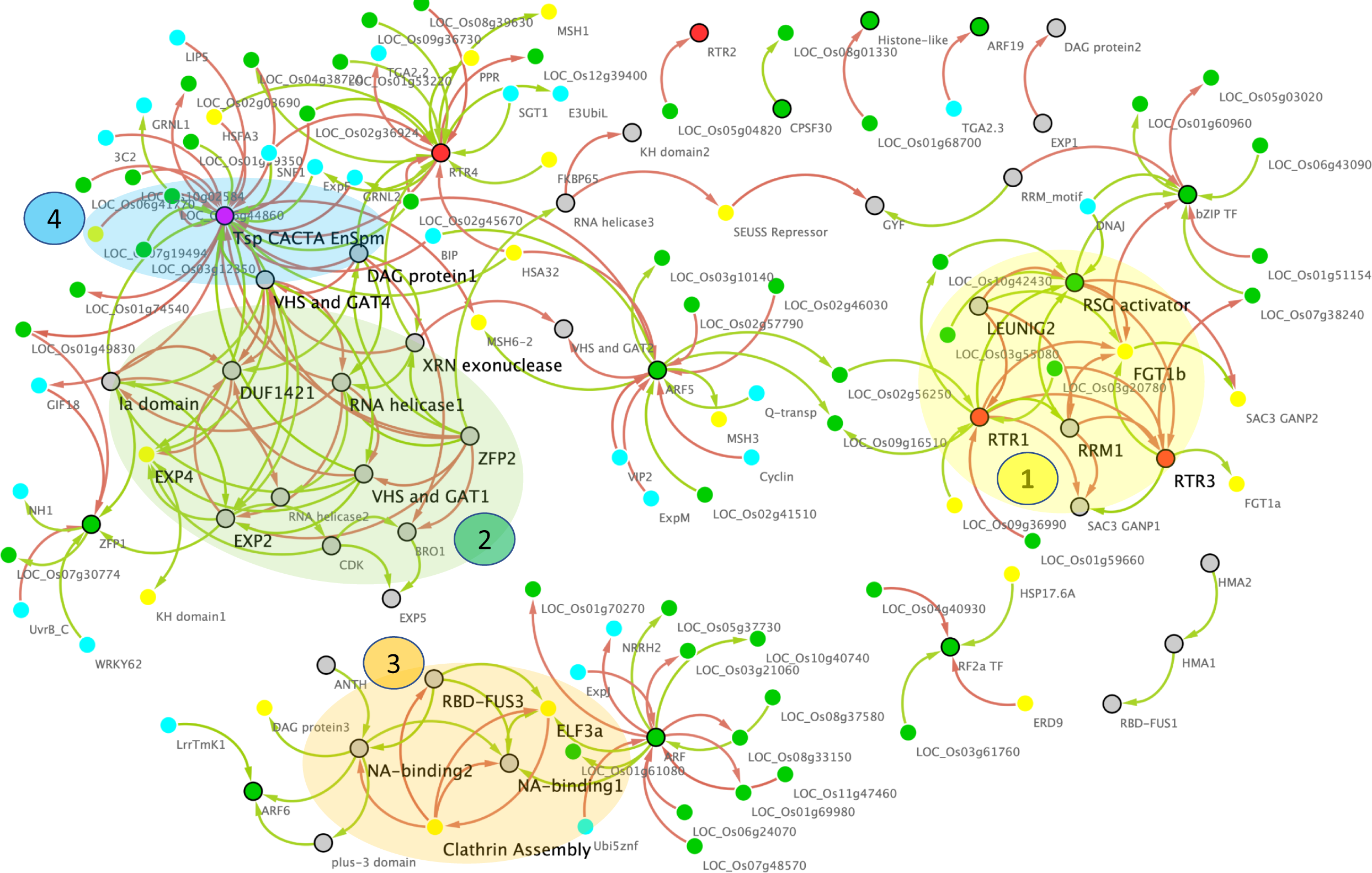
Master gene regulatory network of the Rice Prionome. Nodes represent PrLP genes (black border) and their correlated partners (no border) while edges represent significantly positive (green) or negative (red) expression correlations. Colors of nodes represent Transcription Factors (Green), RTR (Orange) and Transposon CACTA (Magenta). Note the strong overlap of GRN with memory (yellow) and stress (cyan). Four MCODE clusters highlighted in transparent shades, regulatory Hubs in large font.

As can be seen in Figure 7, the network has 208 edges and 139 nodes depicting all 66 PrLPs, including five Ts/RTRs and 11 TFs, along with other significantly correlated TFs, as well as rice genes directly or indirectly implicated in memory and stress events. Most importantly, this network has two large disconnected components, each highlighting the role of rice prionome members as hubs for the currently known data on memory acclimation. Four top ranking MCODE clusters have been highlighted on the network, and each is composed of distinct but tightly inter-connected PrLP genes. Interestingly, only two non-PrLP genes are hubs in these four clusters, and these are homologs of Arabidopsis ELF3 and FORGETTER1 genes, that have very recently been associated heat stress memory via prion-like domains (Jung et al., 2020), and chromatin remodelling mechanism (Brzezinka et al., 2016)respectively. Furthermore, till date, there has been no report connecting these two genes, nor the two above-mentioned memory mechanisms, while the GRN in Figure 7 clearly depicts how pervasively PrLPs act as bridges between the various clusters. For instance, FGT1 lies in the first PrLP cluster where it is closely interacting with RTR1, LEUNIG2, RTR3, RSG activator and RRM1 in the rice prionome, while ELF3 forms part of the third cluster in the GRN of Figure 7, interacting with PrLPs involved in clathrin assembly, NA-binding and RNA binding (RBD-FUS), implying cross-talk between the prion-mediated and epigenetic memory pathways. Other prominent hubs in the PrLP regulatory network are the transcription factors ARF5 and RF2a, as well as Ts/RTRs CACTA and RTR4, earlier observed in the PrLP correlation plot (Figure 6), upregulated in heat stress and female flowers. The four clusters form distinct yet synergistic sub-networks of stress memory within the master regulatory GRN of the prionome, with Transposon CACTA, ARF and RBD-FUS3 having a predominantly antagonistic effect on most memory related genes. For example, transcription factor ARF5 is positively correlated with two rice homologs of the MSH family, that has very recently been implicated in trans-generational heat shock memory (Wierbowski et al., 2020), while it also negatively regulates a homolog of the heat shock protein HSA32 (Charng et al., 2006; Baurle, 2016). The Ts/RTR family member RTR1 is strongly correlated with a rice homolog of a *Chlamydomonas* gene shown to be involved in stress recovery (Hemme et al., 2014), while RTR4 is positively correlated with rice homolog of MSH1, while being inversely correlated with HSA32, closely mimicking ARF5. Similarly, RF2a expression is strongly positively correlated with multiple rice homologs of Hsp17 and inversely correlated with the rice ERD9 gene, reported to be involved in heat stress memory in Arabidopsis (Izadi et al., 2017). In stark contrast, the transposon CACTA is negatively correlated with HSFA3 and MSH family genes, and several heat shock promoter elements, while being positively correlated with HSA32.

Overall, the gene regulatory network of PrLPs, reveals a strongly inter-connected pattern of interaction between TFs, RTRs and genes involved in stress and memory processes, apart from identifying clusters and hubs for future investigation of cross-talk between these molecular factors.

## Discussion

Prions are emerging as one of the most fascinating areas of research in plants, but they have long been documented in animals for a variety of beneficial and detrimental functions; serving as structural modules, components of biofilms, binding to RNA binding and exhibiting enzymatic activity etc (Blanco et al., 2012; Wickner et al., 2015; Wulf et al., 2017; O’Carroll et al., 2020). Their role in memory responses is also widely reported, being causative agents of neurodegenerative diseases associated with memory loss (Prusiner, 2012; Aguzzi and Lakkaraju, 2016; Scheckel and Aguzzi, 2018). In this work we have examined the ‘prionome’ of 39 plant species covering different phyla, from mosses to trees, in order to understand their roles in the plant kingdom.

Prions are well-known to form aggregates or amyloids constituting self-replicating entities. However, formation of amyloid fibrils as a result of protein aggregation is still not shown in plants despite the prediction of PrLPs in their proteomes. Plants are believed to possess lower levels of amyloidosis along with having numerous aggregation-inhibiting compounds, which has indeed been proposed as one of the reasons for longer life span of woody species (Mohammad-Beigi et al., 2019; Surguchov et al., 2019). In this context, we found land plants to possess relatively lower prion densities than the algae, with mosses, as moss genomes having highest prion densities. In algae, these amyloidogenic prion-like proteins may enable biofilm formation as determinants of high mechanical resistance of exo-polysaccharides (Mostaert et al., 2009), also seen in bacteria (Romero et al., 2010), supported further by the fact that we found all four chlorophytes in this study, enriched in signalling-related genes especially, mitogen-activated protein kinases (MAPK), protein kinases, and response regulators. A consensus on overrepresentation of nucleic acid binding functions among the amyloid-forming proteins exists already, and is a common characteristic of PrLPs (Silva and Cordeiro, 2016; Iglesias et al., 2019), and this too, is further corroborated by our results that indicate 40% of PrLPs to have RNA and DNA binding functions in the plant prionomes.

Another interesting category of proteins in the plant prionomes is that of Ts/RTRs, which constitutes 3.4% of the total PrLPs identified from all species. Though prionomes of several organisms have been predicted including those of plants (Antonets and Nizhnikov, 2017) but Ts/RTRs found no mention in any of the previous studies, despite the rice prionome constituting a huge 65% share. Presence of RTRs/Ts-type of PrLPs in plants is intriguing, but indications about their prion-like roles have been reported in human and animal brains, where transposon activity plays a potentially important role in the pathogenesis of prion diseases (Mustafin and Khusnutdinova, 2018). The overwhelming majority of RTR/Ts may also play roles in stress imprinting, as long terminal repeat (LTR) of plants are activated under various stress stimuli, despite being devoid of specific stress response sequences (Alzohairy et al., 2014; Grandbastien, 2015). Reports also suggest heat-induced destabilization of transcriptional gene silencing and activation of particular family of *copia* retrotransposons, named “*Onsen*” (Pecinka et al., 2010; Tittel-Elmer et al., 2010; Ito et al., 2011). Regulation of RTRs has been shown to be mediated by TCP family of TFs playing inhibitory roles in cell division, as positive regulators of aging (Zheng et al., 2018), a phenomenon well-known in humans to be influenced by prionization.

We also found plant prionomes to be enriched in developmental processes like flowering, a phenomenon previously reported (Chakrabortee et al., 2016) where Luminidependens (LD), a protein involved in flowering and its regulation, was reported to mediate plant memory. We also found other flowering-related proteins to possess PrLDs, eg. SEUSS, PFT, FCA and LEUNIG proteins.

We had expected a preponderance of nuclear localization of PrLPs, considering the enrichment of RNA/DNA-binding functions in plant prionomes, but we were surprised to find several PrLPs to exhibit potential dual or even tri-compartment localization as observed from the *in silico* predictions. Of these, RTRs comprise the major fraction. TLS/FUS, an RNA binding PrLP from humans, shows dual localization engaging in nucleo-cytoplasmic shuttling, being primarily localized in the nucleus but under oxidative stress, forming cytoplasmic stress granules (Yamaguchi and Kitajo, 2012). Since prion-like proteins have also been implicated in ‘long term memory’ (Kandel, 2012), they can play a potential role in stress imprinting and memory consolidation, as in case of human Cytoplasmic Polyadenylation Element Binding (CPEB) protein, that enabled persistence of formed memories by transforming in a prion-like manner from a soluble monomeric state to a self-perpetuating translationally active state (Bartsch et al., 1998). In plants, stress memory can be activated upon ‘priming’ with abiotic stress conditions, and can be short term (within a generation) or transgenerational (Slaughter et al., 2012; Kinoshita and Seki, 2014; Wibowo et al., 2016). In this regard, RNA metabolism is a crucial regulatory process, and RNA binding proteins viz. RBDs, KH domain containing proteins or RNA recognition motifs (RRM) are slowly emerging as key regulators of plant responses to environmental constraints (Chinnusamy et al., 2007; Ambrosone et al., 2012; Owttrim, 2013). For instance, RRMs function as chaperones in heat stress (Kang et al., 2013) whereas AtCPSF30 is implicated in redox signaling (Van Ruyskensvelde et al., 2018) and KH domain containing protein is an important upstream regulator for thermotolerance (Guan et al., 2013). We found RNA metabolism related proteins in the plant prionomes and our study also shows the presence of various PrLPs in stress memory and acclimation process. Ten members of the rice prionome were found to be directly or indirectly involved in memory. Two of these are the transcription factors ARF6 and bZIP, while others are LEUNIG1, SUESS repressor, as well as rice homologs of memory associated proteins reported in other plants, such as KH domain1, SAC3/GANP, DAG protein and FCA. Most of these have been reported to be involved in heat stress acclimation but considering the limited information available, it is possible that in future, more PrLPs associated with biotic as well as abiotic stress memory are discovered.

Another mechanism of memory formation in plants is via alterations in chromatin states, such as DNA methylation, histone tail modifications, or paused RNA polymerase II, which can further modify patterns of gene expression that underpin memory responses (Avramova, 2015). In this regard, we found several plant PrLPs to be involved in chromatin remodelling (eg. RNA polymerase II subunits, bZIP, histone methyltransferases etc). In fact, ‘RNA polymerase II’ emerged as an enriched cellular component in the prionomes of several plant species. Further, we find the members of the MSH1 gene family, potential epigenetic sensor of stress (Yang et al., 2020) to be strongly correlated with TF-type PrLPs, suggesting that PrLP-based TFs may act as master regulators of plant memory, and this was explored further through regulatory networks as described later.

The preponderance of DNA binding proteins in the plant prionomes added a new dimension to anticipated roles of PrLPs in plant stress memory consolidation. SWIRM domain containing protein which is a PrLP candidate, has been shown to play a role in temperature induced shaping of epigenetic memory in Norway spruce (Yakovlev et al., 2016). Further, the intracellular transport related SAC3/GANP group of proteins are found to be upregulated during the recovery stage of low temperature stress imposed *P. vulgaris* seedlings (Badowiec and Weidner, 2014). In *Medicago*, the SAC3/GANP family is in fact, associated with ABA upregulation within the co-expression sub-network of seeds from the plants subjected to salinity stress, thereby providing an evidence of transgenerational plasticity (Vu et al., 2015).

The interactome of rice PrLPs further confirmed RNA metabolism related pathways to be the predominant roles, apart from signalling (MAPK pathway). Functional changes induced by the prion form of the cellular prion protein PrP^sc^, involves the p38 MAPK pathway in humans and inhibition of this signalling cascade impedes PrP^Sc^ synaptotoxicity (Fang et al., 2018).

Finally, we have attempted to converge the two current schools of plant memory acclimation mechanisms, namely, chromatin based signals and prion/protein based signals, by conducting a detailed regulatory network analysis of PrLPs that are TFs and RTRs. The availability of high resolution circadian transcriptome of rice combined with available data on TF binding cis-elements, enabled a superimposition of all previously reported plant stress and memory datasets. We generated a high fidelity gene regulatory network for the rice prionome, using available gene expression profile data, filtered by promoter binding site information. Among these interactions, we deciphered an intricate master regulatory network of rice prionome TFs and Ts/RTRs, with at least 50 other rice transcription factors, as well as reported stress and memory pathways. Most importantly, this work revealed four interconnected clusters comprising genes known to be involved in both epigenetic and prion-based signals, involving heat shock memory as well as flowering memory acclimation. The current regulatory network (Figure 7) also has several smaller disconnected clusters of PrLPs, but this may be due to the limited availability of rice prionome gene expression profiles. It must be remembered that all regulatory inferences have been extrapolated from available gene expression profiles of < 25% PrLPs, and yet these domains appear to serve as bridges between epigenetic and prion mediated pathways. We hope that more detailed RNA-Seq experiments in future would fill the gaps and fully connect this network, paving the way for PrLPs as important mediators that converge the two currently known memory mechanisms.

## Conclusion

This work began with an aim to identify and investigate the plant prionome in order to explore a general role in memory, based on previous but limited reports of PrLPs having a role in flowering and plant memory. Plant PrLPs appear as proteins with diverse functions and widespread abundance. Their prion-like functions may be required for either executing their normal developmental roles or in response to environmental cues. Interestingly, we also found a significant enrichment of Ts/RTRs (>60%) in the rice prionome, leading us to wonder whether the epigenetic and protein based memory signals in plants may converge through the prionome, wherein RTRs may have evolved as mediators of memory responses in the plant kingdom. Despite limited data availability, this method enabled the identification of regulatory sub-networks comprising of Ts/RTRs and transcription factors in the prionome, closely connected with genes reported to be involved in stress and epigenetic memory signals, thus intricately connecting the chromatin remodelling and heritable protein pathways for plant memory. In conclusion, our work dissects the possible link between stress and memory in plants which may be executed by mediation of prion-like candidates, thereby helping plants to fortify defences for a stronger or more rapid response in future, by retaining memories of the last event. We hope this will pave the way for more detailed future investigations into precise roles of PrLPs, especially Ts/RTRs with respect to their prion-like properties in plant proteomes.

## Methodology

### Acquisition of proteome sequences and annotation

Proteome sequences of all species used in the study were downloaded from Phytozome (https://phytozome.jgi.doe.gov/) except for *Picea abies* and rice, for which, Congenie database (http://congenie.org/) and Rice Genome Annotation Project (http://rice.plantbiology.msu.edu/) were used, respectively. Annotation for PrLPs above CoreScore ≥ 25 was performed using respective databases.

### Identification of prion-like domain containing proteins

Plant proteomes were analysed for the presence of PrLPs using PLAAC software (http://plaac.wi.mit.edu/). Minimum length for prion domains (L core) was set at 60 and parameter α set at 50. For background frequencies, *Arabidopsis thaliana* proteome was selected. Total number of proteins that contain PrLDs were examined and the proteins having CoreScore ≥ 25 among these were filtered for further analysis. Density was calculated as the ratio of identified PrLPs to the total number of proteins present in the particular species.

### Functional enrichment analysis

Gene Ontology (GO) enrichment was performed using Phytozome (https://phytozome.jgi.doe.gov/pz/portal.html). For detailed analysis of *Oryza sativa* PrLPs, ‘Plant Regulomics’ (http://bioinfo.sibs.ac.cn/plant-regulomics/) and REVIGO (http://revigo.irb.hr/) were used for GO enrichment. Prediction for subcellular localisation of rice PrLPs was carried out using ‘RiceNETDB’ (http://bis.zju.edu.cn/ricenetdb/).

### Tissues-specific, development and stress-based expression analysis

Gene expression analysis of rice PrLPs identified in the study, was carried using Genevestigator (https://genevestigator.com/). Experiments with accession numbers GSE18361, GSE43050, GSE6901, E-MEXP-3718, GSE41647, E-MEXP-2401, GSE14275, GSE6901, GSE58603, GSE19024 were used to extract expression profiles of rice PrLP coding genes under different conditions. Heatmaps depicting gene expression were generated using open-source software R v3.6.1 and R studio v1.2.1335 (R Core Team, 2012).

### Generation of Protein-Protein Interaction (PPI) network

Protein-protein interactors for the 201 rice PrLPs were identified using protein-protein interaction network generation feature in RiceNETDB (http://bis.zju.edu.cn/ricenetdb/). The generated network was then visualized using Cytoscape version 3.7.2 (Shannon et al., 2003).

### Rice genes in stress memory/recovery/acclimatisation

Gene lists for plant stress and memory were mined from literature (Hemme et al., 2014; Brezinka et al., 2016; Georgii et al., 2019; Zuther et al., 2019, Jung et al., 2020); as well as the NCBI GEO database for accessions GSE123072 and GSE112161. For all non-rice genes reported to have been involved in stress or memory responses, we identified the rice homologs by local blast against the rice database, followed by filtering of results by e-value cut-off less than 1e-5, query coverage and identity cut-offs more than 50% and 35% respectively, and selected top ranking gene in rice based on e-value. Both lists and associated references are provided as Supplemental Table 4. Heatmaps were generated using R package.

### Unweighted Gene Co-expression Data Analysis

Time series gene expression data (in Transcripts per million TPM counts) for rice diurnal range across the day-night cycle was downloaded from (Ferrari et al., 2019). Temporal expression patterns for 66 PrLP genes were found in this dataset and extracted using Normalized TPM values (rescaled between 0 and 1 via Min-Max scaling/normalization). Combined parametric and non-parametric (Spearman’s and Pearson’s) correlations were calculated for each pair of genes using R package Hmisc (Harrell Jr and Dupont, 2017). Significantly correlated positive and negative sets of genes were identified by filtering out all correlations below a threshold cut-off of ± 0.8 and P-value < 0.01. The resulting binary expression matrix was used to generate Corrplots for PrLP cluster identification in R corrplot (Wei and Simko, 2019) by applying average linkage clustering to genes and first principle component distance measurements.

### Construction & Clustering of Gene regulatory network

The Gene Regulatory Network for PrLPs was generated from Co-expression Data in a stepwise manner. The first layer of this network included all significant self-self-correlations among PrLPs. The second layer was formed by adding co-expression data for PrLPs and rice TFs. The third and last layer of the GRN was created by adding stress and memory related genes found to be significantly correlated with TFs or RTRs in the rice prionome. Finally, the network was filtered with evidence from cis-binding element information, to retain high fidelity regulatory interactions. For this, promoter sequences of 1700 bp length were extracted from the rice structural annotation (GFF3 format) files for each PrLP. Each such ‘promoter’ comprised of 1500 bp upstream and 200 bp downstream of the annotated transcription start site. Transcription factors that bind to these promoters were identified using Plant transcription regulatory map database (http://plantregmap.cbi.pku.edu.cn/) (Tian et al., 2020). This information was used as supporting evidence to filter the pairs of genes found to be significantly co-expressed in the binary co-expression matrix of PrLPs. The resulting filtered co-expression matrix was exported to an edgelist in SIF format file and visualised using Cytoscape version 3.7.2 (Shannon et al., 2003). MCODE clustering was applied on this network to identify high ranking gene clusters using default parameters (Bader and Hogue, 2003).

## Acknowledgements

G.Y. acknowledges support of NIPGR and funds from BBSRC GCRF Grant ID. BBSRC BB/P027970/1TIGR^2^ESS for this work. C.K. extend thanks to Prof. Ashwani Pareek, Stress Physiology and Molecular Biology Laboratory, School of Life Sciences, JNU, for hosting her during the DST-INSPIRE fellowship and acknowledges DST-INSPIRE grant (IFA-14/LSPA-24) received from the Department of Science and Technology (DST), Government of India. S.G. is supported by a grant from DBT. C is funded by the DBT SRF. S.K.S. thanks SERB for Distinguished Fellowship award

## Author Contributions

Conceptualization, S.K.S & GY; Visualization, C.K., S.G., G.Y.; Investigation, C., S.G., J.P.; Writing-Original draft, S.G., C.K., G.Y.; Writing-Review & Editing-S.K.S, S.L.S-P; Funding, G.Y., C.K., S.K.S.; Supervision, C.K., G.Y.

