## Supplemental Figures for "Plant Prionome maps reveal specific roles of prion-like proteins in stress and memory"

(A)

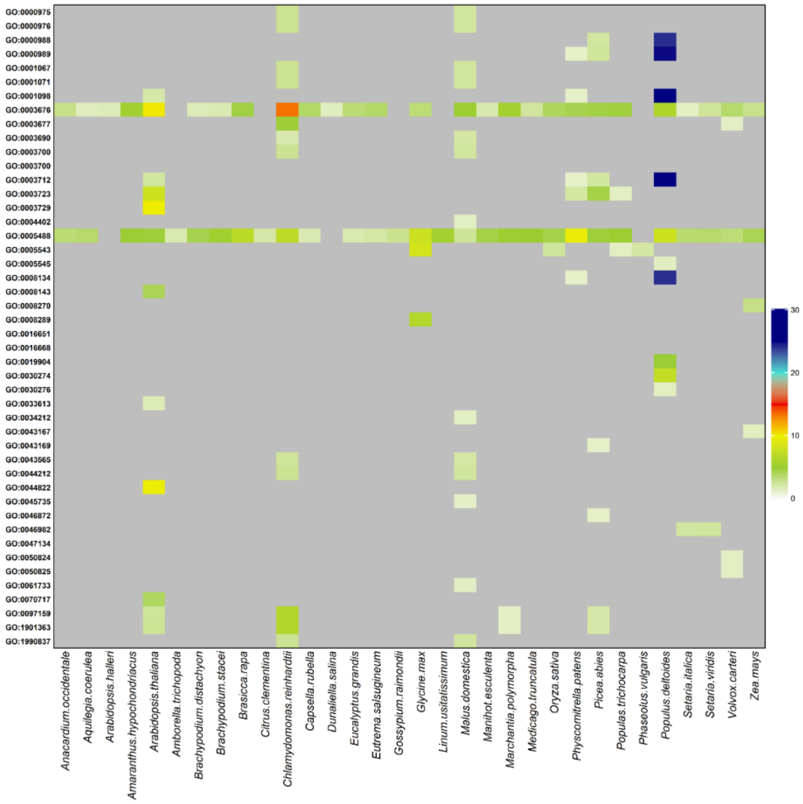
GO:0000975
GO:0000976
GO:0000988
GO:0000989
GO:0001067
GO:0001071
GO:0001098
GO:0003676
GO:0003677
GO:0003690
GO:0003700
GO:0003700
GO:0003712
GO:0003723
GO:0003729
GO:0004402
GO:0005488
GO:0005543
GO:0005545
GO:0008134
GO:0008143
GO:0008270
GO:0008289
GO:0016651
GO:0016668
GO:0019904
GO:0030274
GO:0030276
GO:0033613
GO:0034212
GO:0043167
GO:0043169
GO:0043565
GO:0044212
GO:0044822
GO:0045735
GO:0046872
GO:0046982
GO:0047134
GO:0050824
GO:0050825
GO:0061733
GO:0070717
GO:0097159
GO:1901363
GO:1990837
Anacardium.occidentale
Aquilegia.coerulea
Arabidopsis.halleri
Amaranthus.hypochondriacus
Arabidopsis.thaliana
Amborella.trichopoda
Brachypodium.distachyon
Brachypodium.stacei
Brasicca.rapa
Citrus.clementina
Chlamydomonas.reinhardtii
Capsella.rubella
Dunaliella.salina
Eucalyptus.grandis
Eutrema.salsugineum
Gossypium.raimondii
Glycine.max
Linum.usitatissimum
Malus.domestica
Manihot.esculenta
Marchantia.polymorpha
Medicago.truncatula
Oryza.sativa
Physcomitrella.patens
Picea.abies
Populas.trichocarpa
Phaseolus.vulgaris
Populus.deltoides
Setaria.italica
Setaria.viridis
Volvox.carteri
Zea.mays
30
20
10
0

(B)

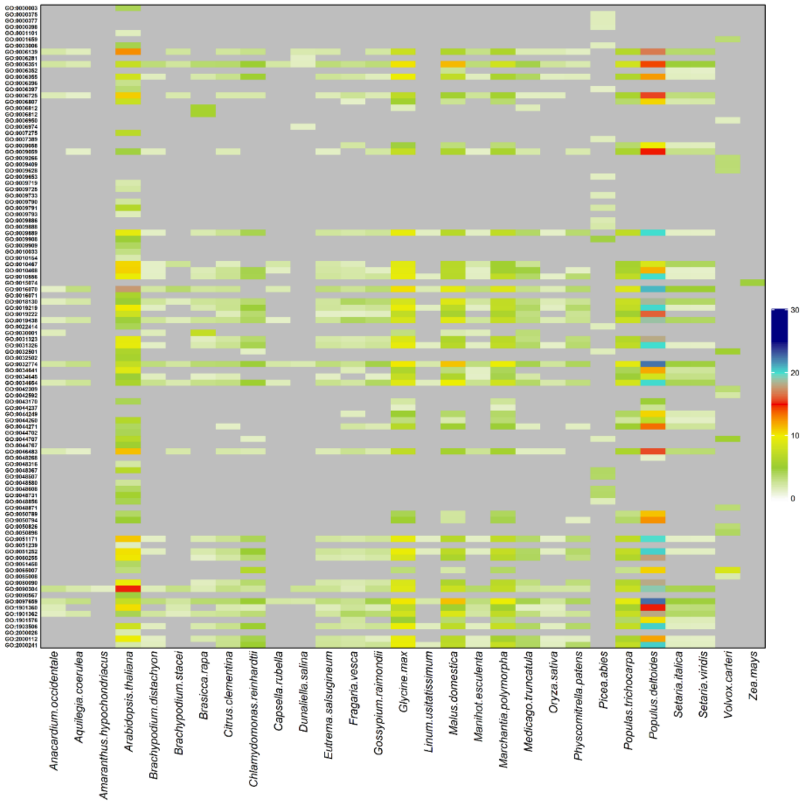
Anacardium.occidentale
Aquilegia.coerulea
Amaranthus.hypochondriacus
Arabidopsis.thaliana
Brachypodium.distachyon
Brachypodium.stacei
Brassica.rapa
Citrus.clementina
Chlamydomonas.reinhardtii
Capsella.rubella
Dunaliella.salina
Eutrema.salsugineum
Fragaria.vesca
Gossypium.raimondii
Glycine.max
Linum.usitatissimum
Malus.domestica
Manihot.esculenta
Marchantia.polymorpha
Medicago.truncatula
Oryza.sativa
Physcomitrella.patens
Picea.abies
Populas.trichocarpa
Populus.deltoides
Setaria.italica
Setaria.viridis
Volvox.carteri
Zea.mays
30
20
10
0

(C)

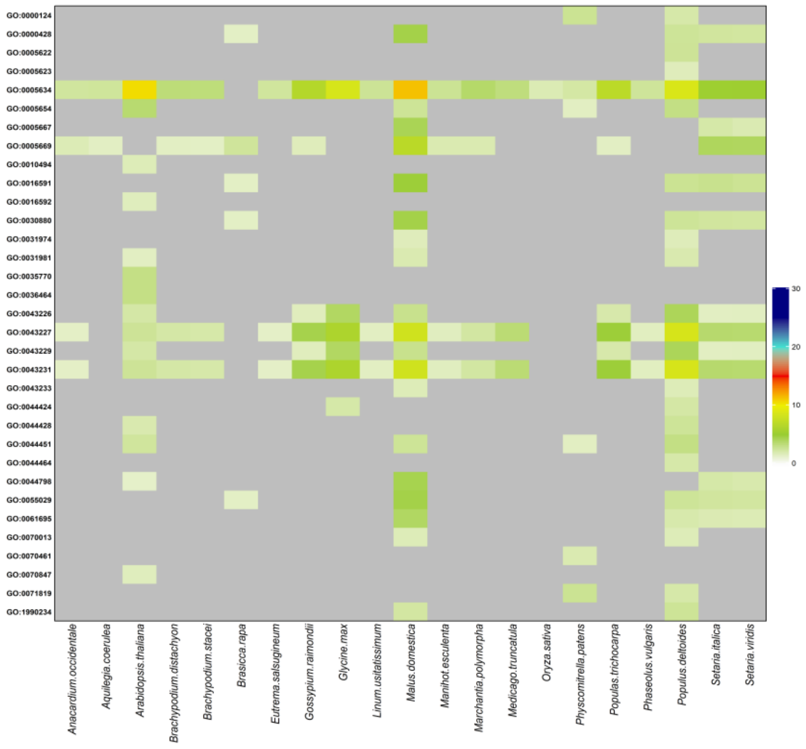


**Supplemental Figure 1.** GO enrichment analysis of PrLPs from organisms used in the study. Heat maps depicting (A) Molecular functions, (B) Biological Processes and (C) Cellular components, enriched in all 39 species. Heatmaps were generated using R package.

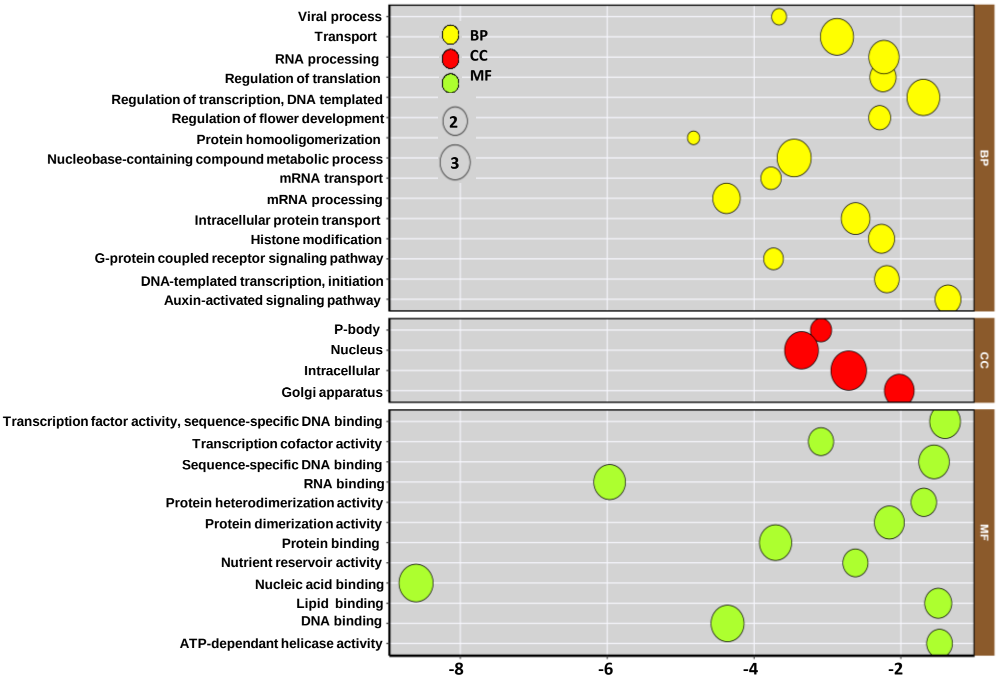


**Supplemental Figure 2.** Gene ontology (GO) enrichment analysis of rice PrLPs. The y axis represents different GO entries belonging to three categories viz biological processes (BP), cellular components (CC) and molecular functions (MF). The x axis represents *p* values (log10 scale) of the unique GO entries. The circle size indicates the frequency of the terms. The more general terms have larger circle area. 'Plant Regulomics' (http://bioinfo.sibs.ac.cn/plant-regulomics/) and REVIGO (http://revigo.irb.hr/) were used for GO enrichment.

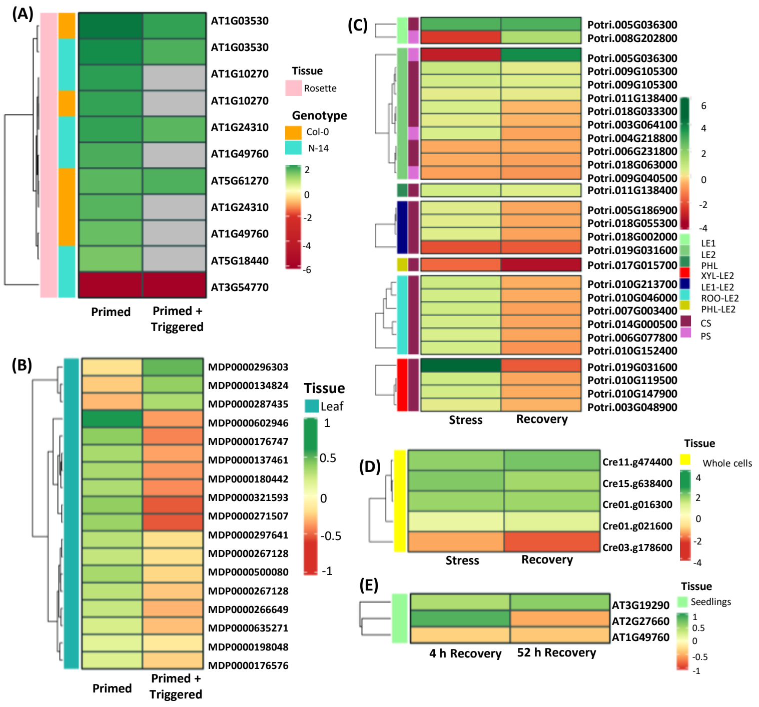


**Supplemental Figure 3**. Plant prionomes involved in stress memory responses. Expression profile of PrLPs in memory responses pertaining to priming as in (A) *Arabidopsis* Col-0 and N-14 genotypes (Zuther et al., 2019), under primed conditions (4°C for 3d followed by 7d of lag phase at 20°C) and subsequent freezing stress (4°C for 3d), and in (B) *M. domestica* (GSE123072), upon treatment with BTH (a salicylic acid analogue) at 14d from propagation and subsequently with flg22 (17d from propagation). PrLP expression profile in memory responses pertaining to recovery phase as in (C) *Populus* spp. (Georgii et al., 2019), in response to periodic (3 intermittent cycles of 6d at 33°C and successively increasing drought from 50%-70% ;2d recovery in between each cycle) or chronic (33°C and gradually increasing drought at 70% for 22d) phase of combined drought-heat stress followed by recovery phase (one week), (D) in *Chlamydomonas* (Hemme et al., 2014), when subjected to heat stress (42°C for 24h) and allowed to recover (25°C for 8h), and in (E) *Arabidopsis* (GSE112161), transcriptome of 4d old seedlings when subjected to heat stress (1h/37°C; 1.5h/23°C; 45 min/44°C) and then allowed to acclimatize at different time points (4h and 52h).

**Supplementary Figure References**


- **Georgii, E., Kugler, K., Pfeifer, M., Vanzo, E., Block, K., Domagalska, M. A., Jud, W., AbdElgawad, H., Asard, H., Reinhardt, R., et al.** (2019). The Systems Architecture of Molecular Memory in Poplar after Abiotic Stress. Plant Cell **31**:346-367.
- **Hemme, D., Veyel, D., Mühlhaus, T., Sommer, F., Jüppner, J., Unger, A.-K., Sandmann, M., Fehrle, I., Schönfelder, S., Steup, M., et al.** (2014). Systems-Wide Analysis of Acclimation Responses to Long-Term Heat Stress and Recovery in the Photosynthetic Model Organism *Chlamydomonas reinhardtii*. Plant Cell **26**:4270-4297.
- **Zuther, E., Schaarschmidt, S., Fischer, A., Erban, A., Pagter, M., Mubeen, U., Giavalisco, P., Kopka, J., Sprenger, H., and Hincha, D. K.** (2019). Molecular signatures associated with increased freezing tolerance due to low temperature memory in *Arabidopsis*. Plant Cell Environ. **42**:854–873.

**Supplemental tables**

**Supplemental Table 1:** Classification of 39 prion-like candidates into 10 different functional categories.

**Supplemental Table 2:** GO enrichment analysis for all the organisms under study.

**Supplemental Table 3:** PrLP candidates predicted in rice prionome with their sub-cellular localisations.

**Supplemental Table 4:** Rice Gene homologs involved in stress and memory.

**Supplemental Table 5**: List of 66 PrLPs with diurnal expression profiles

**Supplemental Table 6:** The Rice Prionome Significant Co-Expression DataSets

**Supplemental Table 7:** Rice Prionome Master Gene regulatory network.

### SUPPLEMENTAL TABLE REFERENCES


- **Baurle, I.** (2016). Plant Heat Adaptation: priming in response to heat stress [version 1; peer review: 2 approved]. *F1000Research* **5**. doi:10.12688/f1000research.7526.1
- **Berry, S., and Dean, C.** (2015). Environmental perception and epigenetic memory: mechanistic insight through FLC. Plant J. **83**:133–148.
- **Brzezinka, K., Altmann, S., Czesnick, H., Nicolas, P., Gorka, M., Benke, E., Kabelitz, T., Jahne, F., Graf, A., Kappel, C., et al.** (2016). *Arabidopsis* FORGETTER1 mediates stress-induced chromatin memory through nucleosome remodeling. *eLife* **5**: e17061.
- **Charng, Y., Liu, H., Liu, N., Hsu, F., and Ko, S.** (2006). Arabidopsis Hsa32, a Novel Heat Shock Protein, Is Essential for Acquired Thermotolerance during Long Recovery after Acclimation. Plant Physiol. **140**:1297-1305.
- **Crisp, P. A., Ganguly, D., Eichten, S. R., Borevitz, J. O., and Pogson, B. J.** (2016). Reconsidering plant memory: Intersections between stress recovery, RNA turnover, and epigenetics. Sci. Adv. **2**:e1501340
- **Izadi, F., Zarrini, H. N., Kiani, G., and Jelodar, N. B.** (2017). Data mining approaches highlighted transcription factors that play role in thermo-priming. Plant Omics **10**:139-145.
- **Jung, J.-H., Barbosa, A. D., Hutin, S., Kumita, J. R., Gao, M., Derwort, D., Silva, C. S., Lai, X., Pierre, E., Geng, F., et al.** (2020). A prion-like domain in ELF3 functions as a thermosensor in *Arabidopsis*. Nature **585**:256–260.
- **Kenchanmane Raju, S. K., Shao, M.-R., Wamboldt, Y., and Mackenzie, S.** (2018). Epigenomic plasticity of *Arabidopsis* msh1 mutants under prolonged cold stress. Plant Direct **2**:e00079.
- **Lämke, J., Brzezinka, K., and Bäurle, I.** (2016). HSFA2 orchestrates transcriptional dynamics after heat stress in *Arabidopsis thaliana*. Transcription **7**:111–114.s
- **Lee, I., Seo, Y.-S., Coltrane, D., Hwang, S., Oh, T., Marcotte, E. M., and Ronald, P. C.** (2011). Genetic dissection of the biotic stress response using a genome-scale gene network for rice. Proc. Natl. Acad. Sci. U S A **108**:18548-18553.

- **Martin, L., Leblanc-Fournier, N., Julien, J.-L., Moulia, B., and Coutand, C.** (2010). Acclimation kinetics of physiological and molecular responses of plants to multiple mechanical loadings. J. Exp. Bot. **61**:2403–2412.
- **Sedaghatmehr, M., Mueller-Roeber, B., and Balazadeh, S.** (2016). The plastid metalloprotease FtsH6 and small heat shock protein HSP21 jointly regulate thermomemory in *Arabidopsis*. Nat. Commun. **7**:12439.
- **Stief, A., Altmann, S., Hoffmann, K., Pant, B. D., Scheible, W.-R., and Bäurle, I.** (2014). *Arabidopsis* miR156 Regulates Tolerance to Recurring Environmental Stress through SPL Transcription Factors. Plant Cell **26**:1792-1807.
- **Suzuki, N., Bajad, S., Shuman, J., Shulaev, V., and Mittler, R.** (2008). The Transcriptional Co-activator MBF1c Is a Key Regulator of Thermotolerance in Arabidopsis thaliana. J. Biol. Chem. **283**:9269–9275.
- **Yang, X., Sanchez, R., Kundariya, H., Maher, T., Dopp, I., Schwegel, R., Virdi, K., Axtell, M. J., and Mackenzie, S. A.** (2020). Segregation of an MSH1 RNAi transgene produces heritable non-genetic memory in association with methylome reprogramming. Nat. Commun. **11**:2214.
